# Expression of the miR-302/367 microRNA cluster is regulated by a conserved long non-coding host gene

**DOI:** 10.1101/497131

**Authors:** Karim Rahimi, Annette C. Füchtbauer, Fardin Fathi, Seyed Javad Mowla, Ernst-Martin Füchtbauer

## Abstract

MicroRNAs (miRNAs) are important regulators of cellular functions. *MiR-302/367* is a polycistronic miRNA cluster including *miR-302*b/c/a/d (collectively termed *miR-302s)* and miR-367. The cluster is located in the intron of a non-coding host gene. *MiR-302s* have been shown to repress mRNAs required for differentiation and to induce pluripotency in somatic cells. The stem cell specific transcription factors OCT4, SOX2 and Nanog drive *miR-302s* expression, however, the reported expression in human and mice indicates a more complex transcriptional regulation. Here we investigate the transcriptional control and the processing of the miR-302 host gene. The murine miR-302 host gene is alternatively spliced, polyadenylated and exported from the nucleus. The regulatory sequences extend at least 2 kb upstream of the transcription start side and contain several conserved binding sites for both transcriptional activators and repressors. Reporter constructs with different upstream regions revealed a significant influence of the more distant regulatory sequences in pluripotent stem cells. The gene structure and regulatory elements like binding sites for activating and repressing transcriptional regulators, splice, and polyadenylation signals are highly conserved between mouse and human. So far, no *miR-302* independent function has been annotated for the miR-302 host gene and we hypothesize that the complex and differential regulation of the miRNA transcription and processing might the reason for its conservation. Thus, regulation or micro-RNA expression might be a so far less recognized function of non-coding RNA genes.

**Author Summary:** Non-coding RNAs constitute a large part of the mammalian genome. Interestingly, some long non-coding RNAs (lncRNA) are transcribed and processed in the same way as mRNAs but lack an open reading frame. Here we give evidence that a so far less recognized function of such lncRNAs could be to supply microRNAs with the complex transcriptional control and processing required for their intricate expression. As an example, we analyzed the regulatory sequences of the miR-302/367 host gene. *MiR-302/367* is a microRNA cluster involved in the regulation of stem cells and cellular differentiation. We show here that the regulatory region is much more complex than anticipated, a complexity that can not be conferred alone by any of the stem cell specific transcription factors which were so far associated with the expression of *miR-302/367*.

## Introduction

MiRNAs are ~22 nucleotide long non-coding RNAs that participate in gene regulation at a post-transcriptional level. They either result in cleavage or translational repression of their target mRNAs. MiRNAs are vital for animal development and compromising their activities is associated with early embryonic lethality or diseases [1–3]. However, both loss and gain of function have been associated with the development and progression of cancer [4]. Dysregulation of miRNAs and their effect on the target genes has been also shown in metabolic disorders [5]. Some miRNAs are known for their potential therapeutic functions [6].

In the genome, miRNAs are localized either intergenic or intragenic, and they can be transcribed as either monocistronic or polycistronic. Intergenic miRNAs are usually transcribed by RNA polymerase III (polIII), while intragenic miRNAs are co-transcribed with their coding or non-coding host genes by RNA polymerase II (polII). If miRNAs are interspersed among Alu repeats, they might be transcribed by either polII or polIII [7–9].

MiRNAs embedded in genes can have feedback effects on their host gene expression [10]. They also have transcriptional regulation activities when associated with the RNA-induced silencing complex (RISC) [3]. It is unknown if there is a principle difference in expression and function between miRNAs located either inter- or intragenic. One might speculate that there is a relation between the complexity of transcriptional regulation and genomic structure. For complex e.g. tissue-specific transcriptional regulation, RNA polIII is not suitable. RNA polII, in contrast, can achieve almost any transcriptional regulation if the appropriate TF binding sites can be recruited as regulatory sequences.

From an evolutionary point of view, it is easily understandable how miRNAs came under the transcriptional control of coding genes. This co-expression allows for negative feedback loops if the host gene at the same time is a target gene. If the host gene is no target gene, the miRNA will gain the same cell or tissue specificity as the host gene. In this line of argumentation, it is, however, less intuitive why miRNAs should be located in non-coding host genes. The two simplest explanations would either be that the host gene has a protein independent function and thus is probably evolutionary older than the embedded miRNA or, that it is solely required to supply the miRNA with a highly regulated polII promoter and a processing scaffold in form of its exon-intron structure and poly adenylation (pA) signal. In this scenario, miRNA and host gene would have evolved together.

The *miR-302/367* (for simplicity here termed *miR-302*) cluster is transcribed from a polII promoter. In human, the *miR-302* cluster is located in the first intron of its host gene on chromosome 4. The hESC-specific expression of the *miR-302* cluster is ascribed to 525 bp immediately upstream of the transcription start site [11, 12]. In both human and mouse, the *miR-302* cluster is located in intron 8 of the LA related protein 7 (*LARP7*) gene, encoded by the opposite DNA strand [11].

In humans, the primary miR-302 host RNA consists of 2-3 exons. *MiR-302* is a polycistronic miRNA cluster including *miR-302*b/c/a/d and miR-367 (with this respective order in the intron). These five miRNAs are generated from the same primary transcript [11]. *MiR-302*a-d are highly related and share the same seed sequence, which is also shared with miR-290-295 in mice and miR-373 in humans. The co-transcribed miR-367 has a different seed sequence [13]. The target genes of miR-367 include *Smad7* and the downstream TGF-ß signaling in cells. It promotes invasion and metastasis of human pancreatic cancer cells [14]. Also, miR-367 causes proliferation and stem cell-like behavior in medulloblastoma cells [15].

The *miR-302s*, on the other hand, have been shown to repress expression of differentiation promoting genes in stem cells, support somatic cell reprogramming [16, 17], and enhance the stemness state of the male germline stem cells [18]. Ectopic *miR-302* expression can mediate induced pluripotent stem cell (iPSC) reprogramming independent of the exogenous addition of transcription factors associated with pluripotency like OCT4, SOX2, KLF4, and MYC [19, 20]. Interestingly, these transcription factors, bind to the promoter region of *miR-302* and tightly regulate murine *miR-302* expression [21] and the expression level of *miR-302* has been reported to correlate with the expression level of Oct4 [16]. Furthermore, the *miR-302* promoter contains three different functional Tcf/Lef binding sites and is a direct target of the WNT/β-catenin signaling pathway. Two of those sites are located within the cluster of OCT4/SOX2/NANOG binding sites, and mutations in the OCT4/NANOG binding sites abolished *miR-302* promoter responsiveness to WNT signaling [22]. Transcription factor COUP-TFII (COUP transcription factor 2) also known as NR2F2 (nuclear receptor subfamily 2, group F, member 2), that can inhibit the transcription of *Oct4* and *Sox2* [23] is translationally repressed by *miR-302*.

In this way, *miR-302s* act as positive regulators of the pluripotency factors OCT4 and SOX2 and stabilize the pluripotency of ESCs [24]. However, as *miR-302s* also target *Oct4*, *Nanog*, and *Sox2*, this has been called an incoherent feedback loop [25]. The functional independence of *miR-302* is further emphasized by the fact that it can regulate tumorigenicity by suppressing both cyclin E-CDK2 and cyclin D-CDK4/6 during G1-S cell cycle transition [26]. For a recent review on *miR-302* function see [27].

During murine embryonic development, expression of the *miR-302* cluster is rapidly down regulated after day 8 of gestation if assayed by whole embryo analysis [13], but little is known about expression in individual poly- or pluripotent stem cells. RT-PCR and whole mount *in situ* hybridization showed *miR-302* expression in the developing lung of murine embryos up to day 15 of gestation. This expression together with the functional studies indicates a strong role of *miR-302* in proliferation and specification of the lung [21].

As *miR-302* is processed from a non-coding gene and as the immediate upstream regulatory region contains a number of stem cell specific transcription factor binding sites, we asked whether the transcriptional regulation of *miR-302* host gene is more complex than the regulation of its main transcription factors like OCT4 or NANOG. This would explain why the *miR-302* cluster is transcribed from its own host gene and not simply is integrated in the intron of one of its main transcription factors.

Our promoter/enhancer analysis revealed a high complexity of the transcriptional regulation of *miR-302*, which exceeds the transcriptional regulation of the individual transcription factors involved. Furthermore, we noticed that the regulatory and RNA processing elements like enhancer, promoter, or splice signals, are highly conserved between human and mouse. This might indicate that transcriptional regulation and processing of the *miR-302* cluster is an important function of the *miR-302* host gene, for which no other function has been annotated so far.

## Results

### Genomic structure of *mmiR-302* cluster and its transcription and processing

In order to analyze the structure and expression of the mmiR-302 host gene, we compared the murine sequence with the annotated *hmiR-302* genomic sequence. In both species, the microRNA cluster is located in the same order and orientation within the intron of a non-coding gene that does not contain any significant open reading frame. In the mouse, none of the potential start codons is preceded by stop codons and none of the potential small peptides can be found in protein databases. We, therefore, consider the miR-302 host RNA to be non-coding. Murine microRNA sequences were obtained from miRBase (www.miRBase.org) [28] and annotated in the intron of the host gene (figure 1). This is comparable to the *hmiR-302* gene [12] with the difference that the human gene contains an alternatively spliced central exon downstream of the *miR-302* cluster [11]. We found no indication of an alternative exon in the murine genome. However, two splice donor sites at the 3’ end of the first exon open the possibility of alternative splicing. Notably, the first splice site is conserved between human and mouse Similar to *hmiR-302*, which has two poly adenylation (pA) signals, *mmiR-302* bears the possibility of alternative polyadenylation as it contains three to four pA signals at the end of the second exon. Figure 1 shows a comparison of the human and murine *miR-302* gene structure.

**Figure 1:**
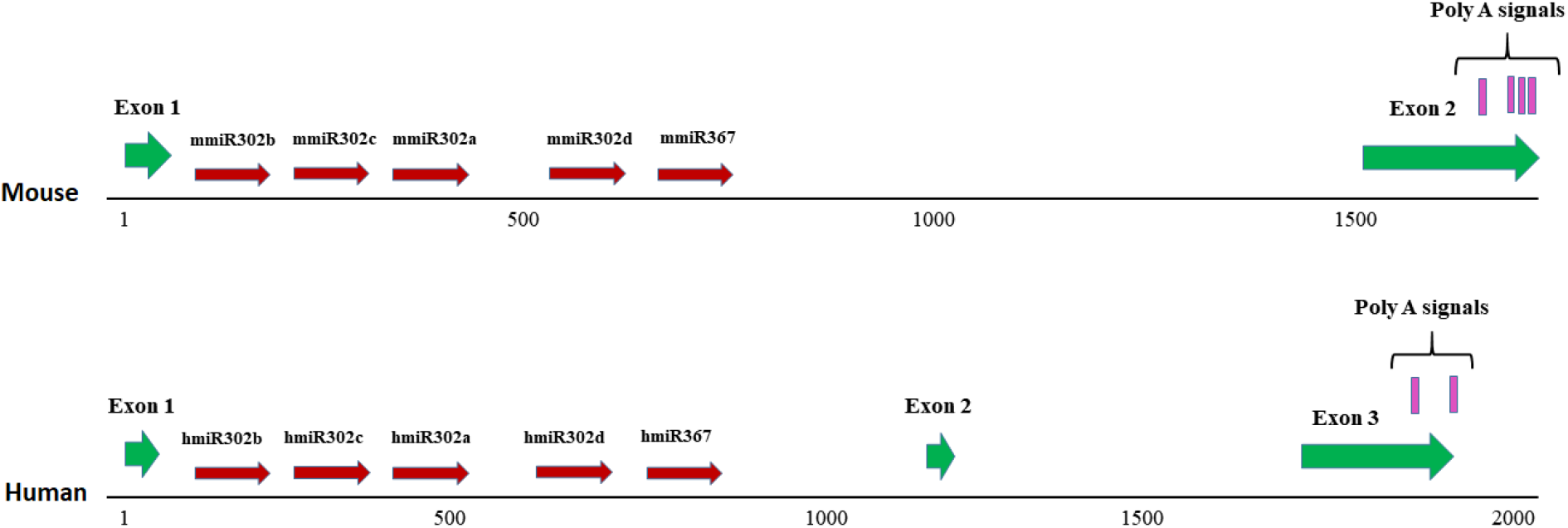
Comparison of the predicted murine *miR-302* gene structure with the homologous human gene. The murine *miR-302* gene (upper part) contains two exons with the microRNA cluster in the intron. The gene is terminated by four potential poly adenylation signals. The human *miR-302* gene (lower part) contains three exons with the microRNA cluster in the first intron. It contains two potential poly adenylation signals at the 3’ end of the third exon, which is slightly shorter than the murine exon 2. Note the different scales.

In order to identify the transcription start site of the *mmiR-302* gene, we performed Rapid Amplification of cDNA Ends (RACE) for the 5’ end of the transcript. Sequencing of the RACE products showed three variations in the transcription start site (variants A, B, and C in figure 2).

**Figure 2:**
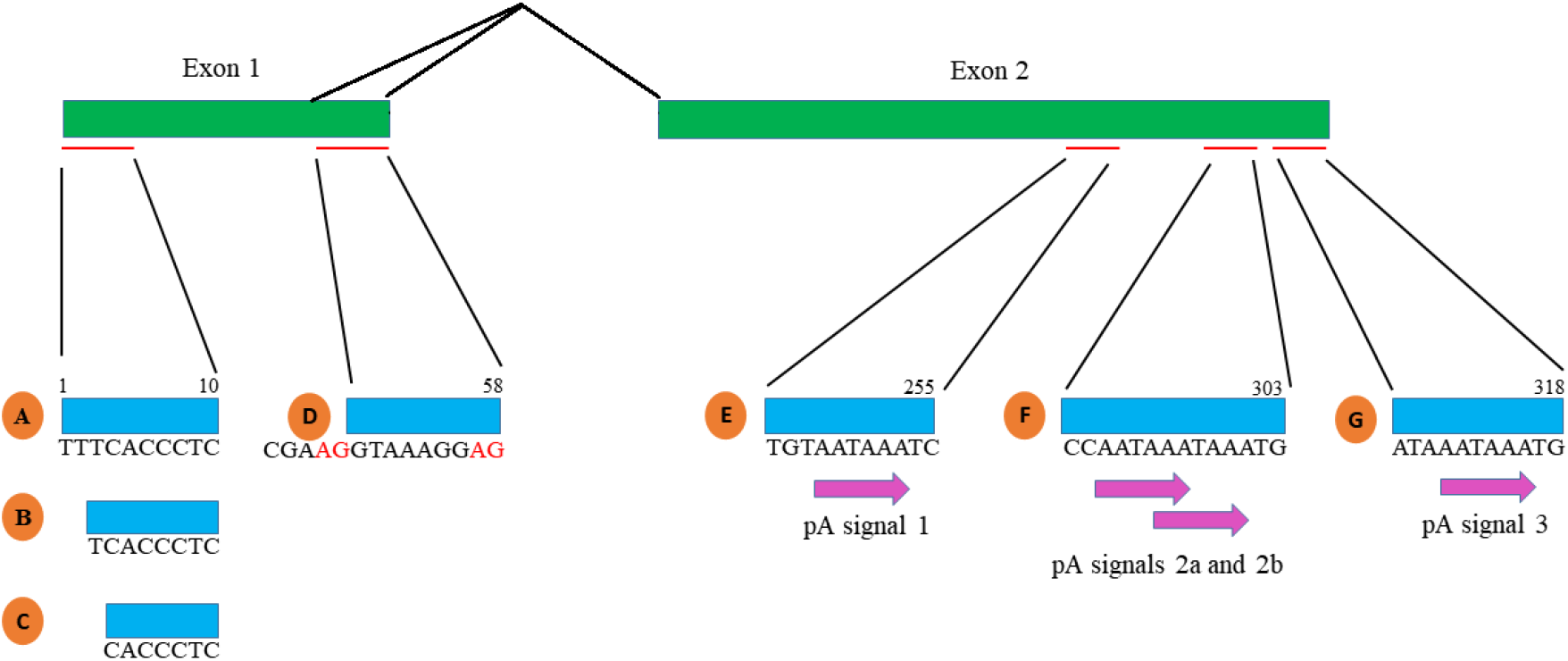
Processing of the murine miR-302 host transcript. The numbering is based on the longest variant of the transcript. Three different transcription start sites (A, B and C) were found. Alternative splicing (D) was found at the end of exon 1, the alternative AG splice donor sites are indicated in red. The RNA was differentially poly-adenylated at three different polyadenylation sites (E, F, and G). All sequences were submitted to GenBank with the following accession numbers: A: KT932380; B: KT932381; C: KT932382; D: KT932383; E: KT932386; F: KT932384; G: KT932385.

RT-PCR using primers located in the first and second exons respectively confirmed both, the absence of a central exon and the alternative use of the two splice donor sites terminating exon 1. Splicing at the second donor site results in a transcript that is 9 bases longer (variant D in figure 2). In three out of three sequences obtained from murine ES cells, this longer version of exon 1 was found while seven out of seven sequences obtained from ES cell-derived teratomas showed splicing at the first splice donor site.

In order to determine which of the potential pA signals, 1, 2a/b, and 3 are used, we performed 3’ RACE on RNA isolated from ES cells and from ES cell-derived teratomas. All six sequences obtained from ES cells were polyadenylated immediately after the last pA signal (pA signal 3 in figure 2) indicating that the overlapping pA signals 2a/b were used which are located 12 and 8 base pairs upstream respectively. Two out of six sequences obtained from murine teratomas used the same pA signal, while the poly-A sequence in three out of six started 12 bases downstream of the last pA signal 3 (G in figure 2). In a single sequence, the first pA signal was used (E in figure 2). The 5’ and 3’ RACE results confirmed that the mmiR-302 host transcript is capped, differentially spliced, and differentially polyadenylated.

### Subcellular localization of the *mmiR-302* spliced host RNA

As the murine miR-302 host RNA is capped, spliced and poly-adenylated, we speculated that it might be exported from the nucleus. To investigate this, we fractionated ES cells and isolated RNA from the cytoplasm (Cyt), endoplasmic reticulum (ER) and nucleus (Nuc) and quantified the miR-302 host RNA relative to *Hprt* mRNA in the respective fractions by qRT-PCR. The efficiency of fractionation was confirmed by Western blot using antibodies against the cytoplasmic protein SEPT9 and tubulin, which is found both in the cytoplasm and the nucleus (figure 3). Even though not 100% pure, the fractions were greatly enriched and showed that the major part of the miR-302 host RNA is exported from the nucleus.

**Figure 3:**
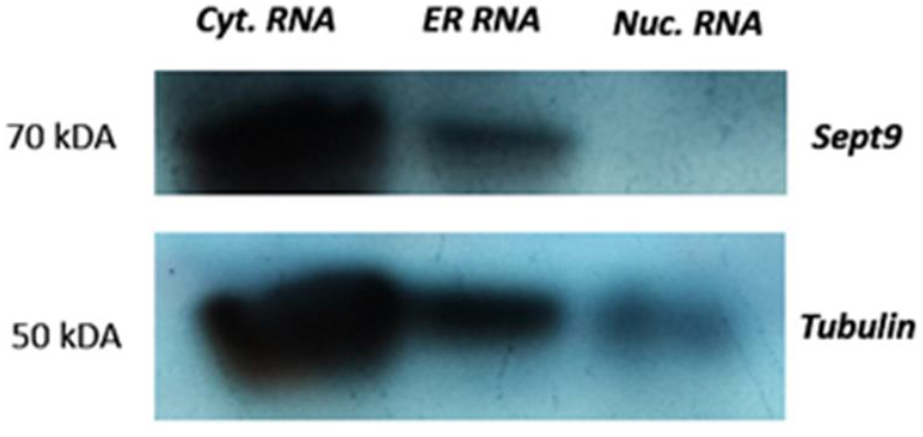
Western blot analysis of sub cellular fractions using Sept9 and tubulin antibodies. The absence of Sept9 in the nucleus shows sufficient separation of the cellular compartments. An equal amount of the fractionation product was used per sample for Western blotting.

The total combined RNA extracted from all three fractions was 83.5 micrograms, of which 71% were from the cytoplasm, 12% from the ER and 17% from the nucleus. Two μg RNA from each fraction, representing 3.33% of the cytoplasmic RNA, 21% of the ER RNA and 14.3% of the nuclear RNA were used for cDNA synthesis. Thus, compared to cytoplasmic RNA, there was a 6.4 and 4.3 fold overrepresentation of ER and nuclear RNA respectively, which is taken into account in figure 4.

For *Hprt* mRNA, Ct values from the cytoplasm (18.5) did not significantly differ from those obtained from whole cell RNA (18.3). The Ct values for the ER and nucleus were much higher (21.7 and 25, respectively), confirming the cytoplasmic localization and efficient nuclear export of *Hprt* mRNA.

**Figure 4:**
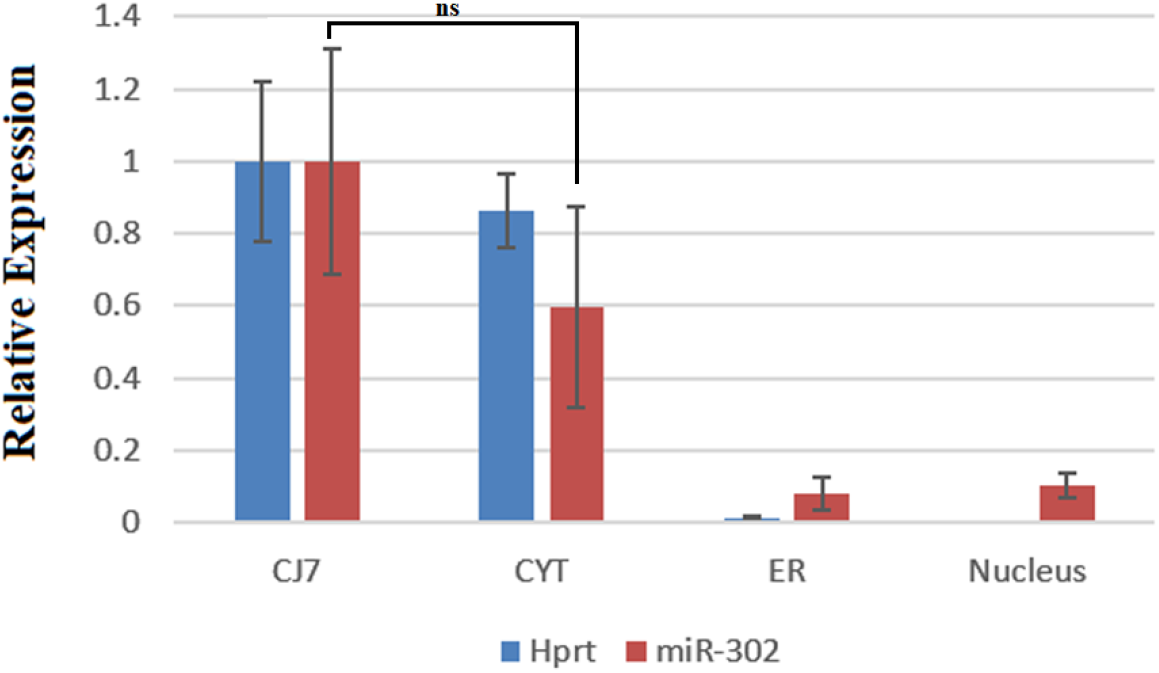
Sub-cellular distribution of mmiR-302 host RNA analyzed by qRT-PCR normalized to the expression of the respective gene in whole ES cells. *Hprt* is used as a control gene, known to be efficiently exported to the cytoplasm. Notably, *Hprt* expression (Ct: 18.29) is almost 30 times higher than the expression of the miR-302 host gene (Ct: 31.76). CJ7: total RNA; CYT: cytoplasmic RNA; ER: RNA associated with the endoplasmic reticulum; Nucleus: nuclear RNA; ns: not significant. Data are presented as mean ± SD, n=3 and *p < 0.05 as significantly level for comparison the samples.

To a lesser degree, the same was true for the mmiR-302 host RNA. The Ct values for all fractions were relatively similar (Cyt: 32.6; ER: 32.8; Nuc: 32.9), but higher than those from total cell RNA (31.8). However, considering the overrepresentation of nuclear RNA used in the assay, these results demonstrate that 70% to 90% of the mmiR-302 host RNA is exported from the nucleus (figure 4).

### Analysis of *miR-302* upstream regulatory sequences

In humans, a 500 bp upstream region has been described as the stem cell specific regulatory element of *miR-302*. This region, which contains among others binding sites for OCT4, SOX2, and NANOG, is highly conserved between human and mouse (figure 5). However, the *miR-302* expression has been reported to be low in naive murine ES cells compared to primed stem cells like murine epiblast stem cells or human ES cells [29]. This is surprising because all the major transcription factors that bind within the first 500 bp in the regulatory sequences located upstream of the transcription start site expected to be responsible for *miR-302* expression are present in all these cells. We, therefore, speculated that the entire promoter/enhancer region should be more complex and possibly contain binding sites for additional TFs including transcriptional repressors. In order to identify such additional regulatory elements, we analyzed 2.1 kb of the upstream genomic sequence using the TRANSFAC, Transcription Factor Binding Sites software (http://www.biobase-international.com/product/transcription-factor-binding-sites) [30].

**Figure 5:**
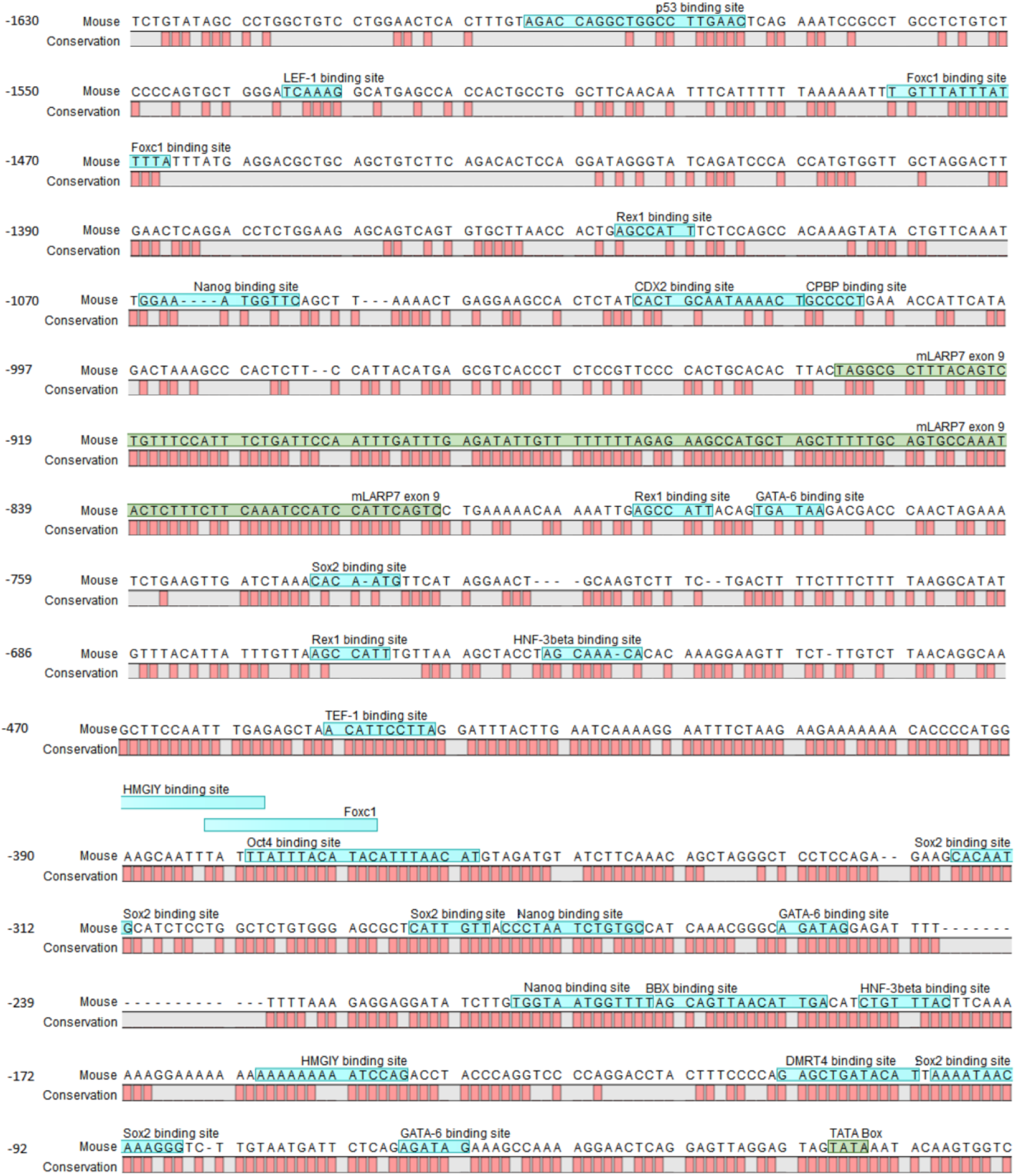
Upstream genomic sequence of murine miR-302 host gene with annotation of some predicted TFs binding sites (blue overlay) and sequence conservation with the human gene (red bars). OCT4, SOX2, and NANOG are the most important stem cell specific TFs that have binding sites in this region. Exon9 of *Larp7* in antisense direction and the miR-302 host gene TATA-box are shown with a green overlay.

Among others, the sequence upstream of −600 contained additional binding sites for OCT4, SOX2, and NANOG, GATA-6, DMRT4, REX1, FOXA2 (HNF-3beta), FOXC1 and TRP53, many of which are conserved between mouse and human (figure 5).

In order to analyze the contribution of the upstream genomic sequences to mmiR-302 host gene expression, we divided the 2.1 kb region into 3 areas which to some degree represent conserved clusters of TF binding sites (figure 6A). We designed luciferase-based reporter vectors containing the different upstream regulatory regions A (+45 to −595), AB (+45 to - 856) and ABC (+45 to −2,120) and used the SV40 promoter upstream of the Renilla luciferase gene as a positive and the promoter-less vector as a negative control. The vector backbone alone drives a considerable ‘background’ transcription of the reporter gene. While this background transcription somewhat reduces the signal strength, it enabled us to monitor activator and repressor function in the same experimental setting.

**Figure 6:**
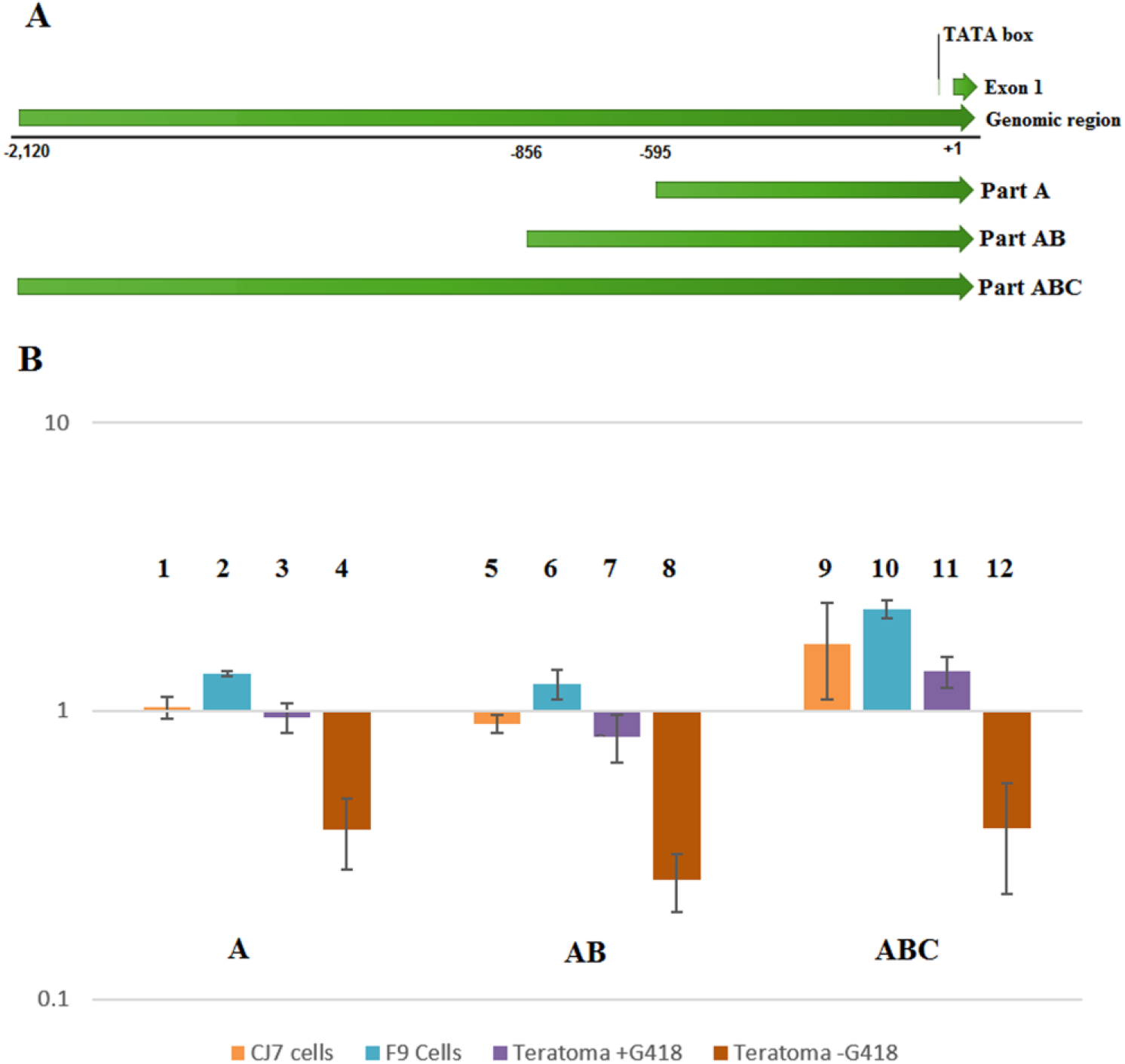
Functional analysis of the 2.1 kb upstream genomic region of the miR-302 host gene. (A) Schematic representation of the 2.1 kb *mmiR-302* upstream genomic region which is divided into 3 parts: A (+45 to −595), B (−596 to −856) and C (−857 to −2,120). (B) Comparison of the promoter/enhancer activity of different regulatory sequences upstream of the *mmiR-302* transcription start site in different cell types. All data are normalized to the expression level obtained from the promoter-less luciferase construct. The baseline expression level is relatively high, which allows to test for repressor activity, which appears as values below 1. F9 cells (lanes 2, 6, 10) show consistently a higher *miR-302* expression than CJ7 ES cells (lanes 1, 5, 9). In teratoma derived cells the expression is lost when selection is stopped (teratoma-G418 lanes 4, 8, 12). Data are shown as mean ± SD, n=3 and *p < 0.05 as significantly level, unless otherwise mentioned. Statistical comparison of data between different cell lines within a single region (A, AB and ABC): part A: 1 vs. 4 p<0.01, 2 vs. 4 p<0.0001 and 3 vs. 4 p<0.01, part B: 5 vs. 8 p<0.01, 6 vs. 8 p<0.0001 and 7 vs. 8 p<0.01 and part C: 9 vs. 10 p<0.05, 9 vs. 12 p<0.0001, 10 vs. 11 p<0.001, 10 vs. 12 p<0.0001 and 11 vs. 12 p<0.0001. The remaining comparisons within one region are not significant. Statistical comparison of data for each cell line between different promoter regions are: 1 vs. 9 p<0.05, 5 vs. 9 p<0.01, 2 vs. 10 p<0.01 and 6 vs. 10 p<0.001. The remaining comparisons of expression level are not significant.

We compared the transcriptional activity of these five reporter constructs in CJ7 ES cells, F9 EC cells and teratoma derived stem cells which were isolated using neomycin resistance driven by the ABC upstream region (Rahimi et al. in preparation).

Only the full-length ABC fragment was able to activate the reporter in all three stem cell like cells, but not in differentiated teratoma-derived cells grown without selection. Compared to ABC, the AB region reduced the backbone driven expression especially in ES and teratoma-derived stem cells (figure 6B). All data were normalized to the baseline expression from the promoter-less vector, therefore values below 1 represent transcriptional repression. Surprisingly, region A, predicted to be the main regulatory region in stem cells, promotes a significant reporter expression only in F9 EC cells. Activation of the miR-302 host gene in CJ7 ES cells and in teratoma derived stem cell like cells required the entire 2.1 kb upstream region. In contrast, but not unexpected, we found that the first 500 bp of the upstream region was sufficient to strongly repressed the baseline expression in differentiated teratoma-derived cells, which had lost their morphological stem cell characteristics.

We observed that expression driven from the 500 bp long A fragment and the 850 bp long AB fragment was lower than from the 2.1 kb long ABC fragment. We therefore searched for potential binding sites of transcriptional repressors located exclusively or predominantly in part AB. We found HBP1 (at −825), SP100 (at −782), BCL−6 (at −419), HIC1 (at −269 and - 848) and GFI1 (at −565) as potential repressors located in this region. Among those HBP1, BCL-6 and HIC1 binding sites are conserved in human and mouse and SP100 and GFI1 belong to few sequences not conserved between human and mouse and only do exist in the mouse. In addition, the distal region of part B (around −850), contains binding sites for XVENT-1, PBX, CDP-CR1 and PIT-1 that are highly conserved among human and mouse.

## Discussion

The structure of the miR-302 host gene is conserved between mice and humans. The main difference is that there is an alternative splice donor site at the end of exon one in the mouse gene while the human homolog contains an alternative second exon not found in the mouse. It is not clear whether the differential splicing and polyadenylation of the miR-302 host gene has any functional consequences for the expression or processing of the embedded microRNAs. However, it is intriguing that both splicing and polyadenylation alternatives are conserved between mouse and human and that our limited data suggest different usage of these sites in different cell types. Both splicing and poly-adenylation have been shown to influence the transcriptional activity of genes and are thus part of the transcriptional regulation [33–35]. It would be interesting to test whether the lnc miR-302 host gene RNA has any function independent of the *miR-302* expression. Unfortunately, the intimate connection of the miR-302 host gene and the *Larp7* gene makes it technically impossible to mutate the miR-302 host gene sequence without simultaneously affecting *Larp7* expression.

Our sequence annotation of the murine and human miR-302 host gene also showed a high conservation of functional elements in the upstream genomic region. Here the majority of transcription factor binding sites and their spatial distribution are conserved. This conservation includes at least 2 kb of upstream genomic sequences, considerably more than the first 500 upstream base pairs which had been annotated in the human gene as *miR-302* promoter [11, 12, 36]. It is noteworthy that the conserved regulatory sequences not only include transcription factor binding sites for supposedly stem cell specific transcription factors like OCT4, SOX2, and NANOG, but also for more tissue-specific transcription factors like GATA-6 or HNF-3ß and even transcriptional inhibitors like HBP1 and SP100. This corresponds well with our analysis of transcriptional regulation of the murine miR-302 host gene and might explain why a supposed stem cell specific micro-RNA is also suggested as a marker for acute heart failure [37].

Different studies have reported the expression of *miR-302s* in ES, EC and iPSC cells [12, 36, 38, 39]. Jesus et al. [12] described that *hmiR-302* is responsible for self-renewal and maintenance of stemness and that its transcriptional regulation depends on stem cell specific TFs. In addition, it has been shown that *miR-302* is one of the factors which maintain pluripotency and affect cell cycle regulation through balancing the expression of Cyclin D1 and D2 [36, 38]. However, it has also been reported that *mmiR-302* expression is elevated in primed stem cells compared to naive ES cells [29, 40], which is more in line with the low level of expression we observed in our experiments. We selected the 2.1 kb upstream region due to the interspecies conservation. Even though the results we obtained are consistent with many published observations on *miR-302* expression, we cannot be sure that our analysis included all regulatory sequences involved in the expression of the miR-302 host gene.

We used the TRANSFAC program to identified binding sites for potential transcriptional repressors in the 5′ upstream region of the miR-302 host gene. The presence of such sites indicates in itself only the possibility of a repressive function. The proof of such function would require ChIP analysis, which so far is not available for the relevant factors and cell types. Alternatively, functional transcription assays might be performed. Assuming, that a transcriptional repression from region AB most likely is mediated by factors binding exclusively or predominantly in this region, we identified HBP1, SP100, BCL-6, HIC1, and GFI1 which are unique for region AB. HBP1 is an inhibitor of the WNT signaling pathway [31] and inhibits the cell cycle [32]. Among the known target genes of HBP1 are *Ccnd1* (encoding Cyclin-D1) and *cMyc* [32], but also *Tcf* and *Lef*, the transcriptional activators of the canonical WNT signaling. Additionally, HBP1 can directly inhibit the DNA binding of the TCF4/β-catenin complex by protein-protein interaction [31]. The murine *miR-302* promoter contains three different functional TCF/LEF binding sites, two of which are located within the cluster of OCT4/SOX2/NANOG binding sites and are essential for WNT/β-catenin-mediated regulation of *miR-302* expression [22]. Thus, HBP1 might inhibit WNT signaling induced miR-302 host gene transcription in three ways: by reducing the expression of TCF and LEF, by inhibiting TCF/LEF DNA binding and by directly repressing miR-302 host gene expression. Since the HBP1 binding site is conserved between human and mice and since both WNT signaling and *miR-302* are involved in stem cell maintenance, it is intriguing to speculate that *miR-302* and the canonical WNT pathway are interconnected in a positive feedback loop that might be negatively controlled by HBP1. SP100 is a potential transcription inhibitor. Interestingly, the SP100 binding site in the regulatory sequence of the miR-302 host gene is only found in mice and not in the human miR-302 host promoter. The latter might indicate a species specific difference in the otherwise highly conserved transcriptional regulation. B-Cell Lymphoma 6 (BCL-6) is a master transcriptional repressor and an oncoprotein which, directly or indirectly inhibits the expression of more than 1000 genes. It is a key regulator gene in multiple myeloma [41, 42]. BCL-6 has also been reported to be an antiviral resistance repressor in follicular T helper cells [43]. Hyper methylated In Cancer 1 (HIC1) is a transcriptional repressor that acts as a tumor suppressor and frequently is inactivated by epigenetic silencing. It has been shown to negatively regulate the WNT signaling pathway [44, 45]. Growth Factor Independence 1 (GFI1) is a zinc finger protein, which acts as a transcriptional repressor. In complex with other factors, GFI1 controls histone modification and affects gene expression. It also has a major effect on regulating self-renewal of hematopoietic stem cells and myeloid and lymphoid differentiation. There is a direct link between AML development and low expression level of *Gfi1* in AML mouse model [46].

In summary, our analysis shows, that region AB has an ambiguous regulatory function, which might explain the cell type specific expression level of the miR-302 host gene in otherwise very similar types of stem cells. It remains to be investigated, to which degree and in which cell types the possible TF binding sites actually are functional.

More than 10,000 of the human non-coding transcripts are longer than 200 bp and are called long non-coding RNAs (lncRNAs) [47]. LncRNAs are usually transcribed by RNA Pol II and share different features with protein coding mRNAs such as capping, splicing, and polyadenylation and often export to the cytoplasm. However, lncRNAs have in average only 2.8 exons compared to 11 exons in mRNAs. In addition, the expression of lncRNAs is on average 10 times lower than the expression of mRNAs [48].

The *miR-302* cluster is embedded in the intron of its host gene, which is transcribed as a lncRNA. This raises the general question why some miRNAs, like *miR-302*, have non-coding host genes. Alternatively, they could be transcribed by RNA pol III or be located in introns of coding genes. Transcription from a RNA pol III promoter does not offer the complex expression pattern observed for many microRNAs. For a microRNA or a microRNA cluster like *miR-302*, an evolutionarily easy way to recruit a pol II promoter would be the localization in an intron of a gene encoding one of its major transcription factors (e.g. OCT4), which should give the appropriate regulation of expression. However, our analysis of miR-302 host gene expression showed that its regulation is far more complex than that of any single transcription factor involved in it.

The intimate sequence relation to the *Larp7* gene excludes functional mutagenesis of the miR-302 host lncRNA, and the absence of function is also theoretically impossible to proof. However, it is conceivable that a major function of the host gene is to provide a regulatory scaffold for the *miR-302* cluster. In this scenario, the export to the cytoplasm might be just a functional by-product of the necessary RNA processing. To our knowledge, this is a new function for genes encoding lncRNAs and might be a more general principle found in a number of RNA pol II transcribed microRNAs that have non-coding host genes.

## Materials and Methods

### Bioinformatics analysis

To investigate the murine *miR-302* gene structure and regulatory sequences and compare it to the human gene, we used the following tools: Ensembl, NCBI and UCSC genome browsers, miRBase (http://www.mirbase.org) [28], miRDB (http://mirdb.org/miRDB) [49], Rfam (http://rfam.xfam.org/) [50], promoter 2.0 prediction server (http://www.cbs.dtu.dk/services/Promoter) [51], Neural Network Promoter Prediction (http://www.fruitfly.org/seq_tools/promoter.html) [52], WebGene (http://www.itb.cnr.it/webgene) [53], functional RNA database (http://www.ncrna.org/frnadb) [54], GeneMark (http://exon.gatech.edu) [55], Transcriptional Regulatory Element Database (http://rulai.cshl.edu/cgi-bin/TRED/tred.cgi?process=home) [56], Mammalian Promoter Database (http://rulai.cshl.edu/cshlmpd/release.html) [57], Mouse Genome Informatics (http://www.informatics.jax.org) [58], CLC main work bench (https://www.qiagenbioinformatics.com/products/clc-main-workbench) and Transcription Factors Binding Sites tool, TRANSFAC (http://www.biobase-international.com/product/transcription-factor-binding-sites) [30], MaxEntScan (http://genes.mit.edu/burgelab/maxent/Xmaxentscan_scoreseq.html), GeneSplicer (http://ccb.jhu.edu/software/genesplicer) [59] and SplicePort (http://spliceport.cbcb.umd.edu) [60].

### Statistical data analysis

Analysis of Variance (ANOVA) was used to determine whether the different expression levels among the genes and different promoter elements are statistically significant. In the all experiments triplicates were analyzed and the results are shown as the mean ± standard deviation (SD). The value of *p < 0.05 was considered to be statistically significant. If not otherwise mentioned, the significance of the difference between the samples was analyzed by the Tukey test, as a Post Hoc test. All the statistical data analyzing were performed using GraphPad Prism software version 7.00 for windows, “www.graphpad.com”.

### DNA and RNA preparation

Genomic mouse DNA was isolated according to [61]. GeneJET Gel Extraction Kit (Thermo Scientific) was used for gel purification. Total RNA was purified using the TRIzol (Invitrogen).

### PCR, RT-PCR and qRT-PCR reactions mix

AmpliTaq Gold 360 Master Mix (ThermoFisher) or Pfu DNA polymerase (Fermentas) were used for PCR and RT-PCR reactions. cDNA was synthesized from 2μg RNA using the M-MLV Reverse Transcriptase (Invitrogen) in a 20 μl reaction mix, 2 % of this was used as template in the qRT-PCR reactions using Platinum SYBR Green qPCR SuperMix-UDG (Invitrogen). Primer sequences and PCR programs are described in the supplement.

### Rapid Amplification of cDNA Ends (RACE)

Rapid Amplification of cDNA Ends (RACE) for the 5’ and 3’ ends was performed using ExactSTART™ Eukaryotic mRNA 5′- & 3′-RACE Kit (Epicentre). For 5’-RACE “mmiR-302 t Rev4” was used as reverse and internal primer, for 3’-RACE “mmiR-302 t fwd6” was used as forward and internal primer. Primer sequences and PCR programs are described in the supplement.

### Cloning protocols

All subcloning was performed using standard molecular techniques. TOPO^®^ TA Cloning (Invitrogen) was used to clone PCR products. All sequencing was performed by GATC Biotech, Konstanz, Germany. Sequences were analyzed using CLC main workbench (Qiagen).

### Luciferase assay vectors

Different upstream genomic regions of the miR-302 host gene were inserted into the psiCheck2 Promega vector (GenBank Accession Number: AY535007) upstream of the *Renilla* luciferase gene replacing the SV40 promoter. *Firefly* luciferase driven by the TK promoter was used as an internal reference gene on the same vector. The vector with and without the SV40 promoter served as positive and negative control respectively. FuGene6 (Promega) transfection reagent was used for all transient transfections. Dual-Luciferase^®^ Reporter Assay System kit (Promega,) was used for luciferase assays.

### Teratoma derived cells

Teratomas were generated by subcutaneous injection of 50 μl Hank’s solution containing 1600 CJ7 ES cells stably transfected with a neomycin resistance gene driven by 2.1 kb upstream genomic region of the mmiR-302 host gene (part ABC). Two 7 month old male 129Sv/Pas mice were injected on both sides of the back without anesthesia. Mice were kept under standard conditions with food and water supplied ad libitum, monitored daily and sacrificed by cervical dislocation after 21 days when the tumors had reached a size of approximately 1 cm^3^. Tumors were carefully dissected avoiding contamination with surrounding tissue and divided for RNA preparation, histology, and cell culture. For cell culture, samples were cut into small pieces of 3-4 mm, washed 3 times with calcium and magnesium free PBS and immersed in 0.25% trypsin for 6 hours at 4 degrees. Then the excess trypsin was removed and the tissue was incubated for 30 minutes at 37 degrees. Trypsin was inactivated with DMEM containing 10% FBS and, 10^6^ cells were seeded per well of 12 wells plate in ES cell medium without selection. After two weeks plates were confluent with differentiated cells like myoblasts and adipoblasts. No stem cell like cells were visible. At this time point, 300 μg/ml G418 was added to the medium, which efficiently removed the differentiated cells. After a few days, ES cell like cells appeared in the culture. In some cases selection with G418 was stopped 4 days prior to the transfection of the luciferase reporter constructs.

### Cell fractionation

Trypsinized and washed cells from a confluent 6 cm dish were lysed in 500 μl fractionation buffer (10 mM Tris-HCl pH 7.4, 10 mM NaCl, 2.5 mM MgCl_2_, 200 μg/ml digitonin) by gentle pipetting. After 10 minutes of gentle rotation at 4 °C the lysates were centrifuged at ~1000 g for 5 minutes at 4 °C in order to pellet the nuclei. The supernatant was then centrifuged at 18000 g at 4 °C for 30 minutes. The new supernatant represented the cytoplasmic (Cyt) fraction. The nuclei pellet was re-suspended in 500 μl homogenization buffer (100 mM Tris-Cl pH 7.5, 150 mM NaCl, 600 mM KCl, 147 mM sucrose, 0.3% NP40) and was centrifuged again at ~1000 g for 5 minutes at 4 °C to pellet the nuclei. The supernatant was then centrifuged at 18000 g for 30 minutes at 4 °C, the supernatant of this centrifugation was assumed to represent the endoplasmic reticulum (ER) fraction. The nuclei pellet was re-suspended in 500 ul S1 solution (250 mM sucrose, 10 mM MgCl_2_) and added on top of an equal volume of S2 solution (880 mM sucrose, 0.5 mM MgCl_2_) and centrifuged at 2800 g for 10 minutes at 4 °C. The supernatant was discarded and nuclei pellet was resuspended in 500 ul wash buffer (10 mM Tris-Cl pH 7.5, 15 mM NaCl, 60 mM KCl) and spined at 1000 g for 5 minutes at 4 °C. The pellet was again resuspended in 500 ul PBS and centrifuged at 1000 g for 5 minutes at 4 °C. The pellet represented the nuclear (nuc) fraction. For RNA preparation, 1 ml Trizol was added to each fractionation.

### SDS-PAGE and Western Blot

For Western blot, 10μl from each cell fraction were mixed in 10μl Laemmli buffer (S3401-10VL Sigma-Aldrich) and separated on a 10% acrylamid gel containing SDS (3.33 ml protogel, 2.5 Tris pH 8 1.5M, 50μl SDS, 10μl TEMED and 100μl Aps) using a Novex Western blot chamber. Protein bands transferred onto Immobilo-FL Transfer Membrane. To block unspecific protein binding, the membrane was incubated in 1X TBS-T buffer supplemented with 5% skimmed milk and 0.1% Tween-20 for 1 hour at room temperature. Primary antibodies incubation was carried out overnight at 4°C followed by secondary antibody incubation at room temperature for 1 hour. Signals were detected using ECL Select Western Blotting Detection Reagent (RPN2235 GE Healthcare) according to the manufacture’s manual. Both primary antibodies, rabbit anti Sept9 (10769-1-AP Sept9 polyclonal antibody from Protein-Tech) and mouse anti alpha tubulin (600-401-880 mouse monoclonal antibody from RockLand) were diluted 1:1000 in TBS-T buffer supplemented with 5% skimmed milk and in dilution. Secondary antibodies, Anti-Rabbit IgG Peroxidase antibody produced in goat (A6154 from Sigma-Aldrich) 1:7500 and rabbit anti mouse HPR antibody from Dako (P0260) 1:5000, were diluted in 1X TBS-T buffer supplemented with 3% skimmed milk.

### Ethics Statement

All animal experiments have been performed with permission and according to the regulations of the Danish Animal Experiments Inspectorate, the legal authority under the Danish Ministry of Environment and Food.

## Supporting information

## List of abbreviations

ES cells: Embryonic stem cells
EC cells: Embryonic carcinoma cells
iPSCs: Induced pluripotent stem cells
miRNAs: MicroRNAs
polII: RNA polymerase II
polIII: RNA polymerase III
RISC: RNA-induced silencing complex
TFs: Transcription factors
RACE: Rapid Amplification of cDNA Ends
LncRNAs: Long non-coding RNAs

## Supporting Information Legends (supplements)

Supporting Information 1: Table S1: Primer sequences

Supporting Information 2: Programs used for PCRs

