## Supplementary material for "Expression of the miR-302/367 microRNA cluster is regulated by a conserved long non-coding host gene"

**Supplements:**

**List of primer sequences**

Primers were designed using CLC main work bench (Qiagen) and checked by NCBI primer-BLAST and ordered from Sigma-Aldrich and used at a concentration of 1 µM for PCR..

| Name | 5′-3′ sequence | bp |
|---|---|---|
| mmiR-302 promoter Fwd | AAGAATATTAATGTTTCCTGGTTGCTTCTAAT | 32 |
| mmiR-302promoter Rev exon 1 | AAGAATGCTAGCTGACCGCCTCCCAAAGAGTCCTGTTC | 38 |
| mmiR-302 transcript fwd1 | CTCCGAGGACAGAACAGGA | 19 |
| mmiR-302 transcript fwd2 | AGAACAGGACTCTTTGGGAG | 20 |
| mmiR-302 transcript fwd3 | TCACGAAGGGTCCCACA | 17 |
| mmiR-302 transcript fwd4 | CACGAAGGTAAAGGAGGGT | 19 |
| mmiR-302 transcript fwd5 | GCTTTTGCTGCTTGCTCTTTT | 21 |
| mmiR-302 transcript fwd6 | AACCACATTGCCACATTTCCCA | 22 |
| mmiR-302 transcript rev1 | ATTTATAATTCATTTATTGG | 20 |
| mmiR-302 transcript rev2 | TATTACAGGAGTTGCTC | 17 |
| mmiR-302 transcript rev3 | GCTCCCCAAAAATGTTACTCA | 21 |
| mmiR-302 transcript rev4 | GGATTTGCCTTTGTGGAA | 22 |
| adapter overhang primer | GCGAGCTCCGCGGCCGCGTTTTTTTTTTTT | 30 |
| anchor overhang primer | GGCCACGCGTCGACTAGTACTTTTTTTTTTTTTTTTT | 37 |
| mmiR-302pFwd1 | AAGAATACCGGTCTGGAGTTGCTTTGTTTTC | 31 |
| mmiR-302pRev1&2 | AAGAATCATGTTAAAGCAGAGGGGA | 25 |
| PCR primer 1 (PP1) | TCATACACATACGATTTAGGTGACACTATAGAGCGGCCGCCTGCAGGAAA | 50 |
| PCR primer 2 (PP2) | TAGACTTAGAAATTAATACGACTCACTATAGGCGCGCCACCG | 42 |
| HPRT +1 | GCAAGCTTGCTGGTGAAAAGGA | 22 |
| HPRT -1 | GCAGAUGGCCACAGGACUAGAACA | 24 |

Table S1: Primer sequences

### Programs used for PCRs

***mmiR-302* promoter (short region A) PCR:**

94 ˚C 5 min, (94 ˚C 1 min, 58 ˚C 45 sec, 72 ˚C 1 min) X 35, 72 ˚C 7 min

Forward primer: mmiR-302 promoter Fwd1

Reverse primer: mmiR-302promoter Rev exon1

***mmiR-302* spliced host RNA PCR:**

94 ˚C 5 min, (94 ˚C 1 min, 56 ˚C 1 min, 72 ˚C 1 min) X 35, 72 ˚C 7 min

Forward primer: mmiR-302 transcript fwd2

Reverse primer: mmiR-302 transcript rev3

***Hprt* PCR:**

94 ˚C 5 min, (94 ˚C 45 sec, 60 ˚C 45 sec, 72 ˚C 1 min) X 30, 72 ˚C 7 min

Forward primer: HPRT +1

Reverse primer: HPRT -1

**5’ RACE PCR:**

94 ˚C 5 min, (94 ˚C 1 min, 58 ˚C 1 min, 72 ˚C 1 min) X 35, 72 ˚C 7 min

Forward primer: PCR primer 1 (PP1)

Reverse primer: mmiR-302 transcript rev4

**3’ RACE PCR:**

94 ˚C 5 min, (94 ˚C 1 min, 54 ˚C 1 min, 72 ˚C 1 min) X 35, 72 ˚C 7 min

Forward primer: mmiR-302 transcript fwd6

Reverse primer: PCR primer 2 (PP2)

**Program for all qPCRs:**

94 ˚C 10 min, (94 ˚C 30 sec, 56 ˚C 30 sec, 72 ˚C 30 sec) X 40, 95 ˚C 1 min, 55 ˚C 30 sec, 95 ˚C 30 sec
